# Optimized reference region and the effect on test-retest reliability and detection of Parkinson’s disease with [^11^C]UCB-J

**DOI:** 10.1101/2025.10.15.682460

**Authors:** Nikkita Khattar, David Matuskey, Jean-Dominique Gallezot, Mika Naganawa, Sophie E. Holmes, Faranak Ebrahimian Sadabad, Mahsa Mayeli, Irina Esterlis, Christopher H van Dyck, Adam P Mecca, Deepak Cyril D’Souza, Nabeel B. Nabulsi, Sjoerd J. Finnema, Yiyun Huang, Richard E. Carson, Takuya Toyonaga

## Abstract

[^11^C]UCB-J is a radioligand targeting synaptic vesicle glycoprotein 2A, used to image synaptic density. For quantification, a small-volume centrum semiovale area was previously optimized as a [^11^C]UCB-J reference region (CS2mL); however, its high variability resulted in reduced reliability. Herin, we evaluated an alternative reference region method to assess longitudinal test-retest reliability and detection of Parkinson’s disease (PD). For estimating distribution volume ratio (DVR), CS2mL and eleven white matter (WM) reference regions (range: 0.5-200 mL) were generated using the Freesurfer WM map. Same-day and longitudinal test-retest variability (TRV) were assessed (24 healthy subjects (HS); n=10 same-day and n=20 longitudinal HRRT scans, range: 7-1028 days). Each reference region was used to evaluate the substantia nigra (SN) and caudate DVRs in HS (n=25) and PD (n=20). The 10mL WM reference region yielded [^11^C]UCB-J DVR measurements with reduced variability in TRV (same-day: 10mL: 1.2±5.7%, same-day: CS2mL: -0.9±9.2% longitudinal: 10mL: 1.5±7.0%, CS2mL: 1.6±11.9%,) while maintaining <10% volume of distribution difference, compared to CS2mL. Further, a significant difference between PD and HS groups in SN and caudate DVRs was found using 10mL, with greater effect size (Cohen’s d 0.61 for SN and 0.66 for caudate) compared to CS2mL (0.38 for SN and 0.43 for caudate).

## Introduction

Synaptic vesicle glycoprotein 2A (SV2A) is homogenously expressed on pre-synaptic vesicles of neurons throughout the central nervous system (1, 2). Measurements of SV2A are useful as a surrogate marker for synaptic density, and recent advancements in positron emission tomography (PET) imaging have rendered *in-vivo* synaptic density measurements possible (2–5).

[^11^C]UCB-J, a radioligand targeting SV2A, has exhibited high specific binding, excellent brain uptake, and fast kinetic characteristics (5, 6). As such, [^11^C]UCB-J PET has been applied extensively in humans to show differences in synaptic density in several neurological disorders including Alzheimer’s disease (AD) (7, 8) and Parkinson’s disease (PD) (9, 10).

To validate the use of [^11^C]UCB-J in clinical studies, our group performed a kinetic evaluation of [^11^C]UCB-J in healthy subjects (HS) and we reported excellent same-day test-retest reproducibility of [^11^C]UCB-J (6). Nevertheless, high longitudinal reliability is critical, particularly when studying neurodegenerative diseases that are progressive over several years, to evaluating pathological changes in synaptic density with time (11, 12). Thus, optimizing quantification methodology can enhance statistical power to detect within-subject changes, allowing for clinical trials to have smaller population sizes or shorter duration.

Further, a reference region with minimal displaceable activity is ideally necessary to fully quantify and evaluate the specific binding of [^11^C]UCB-J. Binding potentials (*BP*_ND_) have been derived using the centrum semiovale (CS) – a white matter (WM) region-of-interest (ROI) that has been optimized to have minimal displaceable activity and spill-in from gray matter (GM) regions (5, 7, 13) as defined by the small-volume version of the CS by Rossano et. al. (referred to as CS2mL) (13). ROIs derived from the WM can be appropriate reference regions due to the low uptake of SV2A compared to the GM (5). However, due to its small size (< 2 mL), the CS2mL ROI introduces high variability into the measurements of specific binding, and longitudinal reliability was not considered in the initial selection of this CS ROI size. Mecca et al proposed using the whole cerebellum as a reference region for evaluating differences in distribution volume ratios (DVRs) between AD patients and HS because the cerebellum exhibits a minimal difference in specific binding between these groups, and remains relatively intact in AD (7). Although the cerebellum provides a useful and appropriate reference ROI for studies of diseases wherein it is spared, cerebellar pathology is implicated in many neurodegenerative diseases (14, 15). To this end, a WM ROI with less noise and minimal specific binding could be a useful reference region for many disorders.

Thus, to reevaluate reference regions for [^11^C]UCB-J, we created multiple sized WM ROIs for reference regions and performed same-day and longitudinal evaluations of test-retest reliability in HS. To optimize the WM reference ROI size, we aimed to balance the high variability introduced using a small ROI with the bias induced by a larger ROI. As such, we compared the volume of distribution (*V*_T_) values among each reference ROI as well as the SD of the DVR test-retest variabilities (TRVs) computed using each reference region. Finally, we utilized the optimized reference ROI to reanalyze a subset of [^11^C]UCB-J PET images from a previous cross-sectional study of PD patients and HS (10), to evaluate the reference region method when applied toward studying human pathophysiology.

## Material and Methods

### Human participants

Twenty-four HS participated in the test-retest study (age: 40.3±14.3 years, M:F = 13:11, 10 pairs of same-day scans, 20 pairs of longitudinal scans). These data are an extension of a previous same-day test-retest study (6). A subset of subjects studied in Holmes and Honhar et. al (10) who had arterial blood sampling were included in the PD cohort: twenty-five HSs (age: 60.6±8.8 y, M:F = 11:14) and twenty PD subjects (age: 61.6±8.6 yr, M:F=9:11, Movement Disorders Society-Unified Parkinson’s Disease Rating Scale III: 30.5±9.0, Hoehn & Yahr: 2.0±0.0, Montreal Cognitive Assessment: 26.4±2.3, Disease duration: 4.7±3.0). The data were collected from several protocols approved by the Yale University Human Investigation Committee and the Yale University Radiation Safety Committee, and in accordance with the United States federal policy for the protection of human research subjects contained in Title 45 Part 46 of the Code of Federal Regulations (45 CFR 46). Written informed consent was obtained from all participants after complete explanation of study procedures.

### Injected Radiotracer

[^11^C]UCB-J was synthesized according to previously described procedures (16). The injected radioactivity of [^11^C]UCB-J was 491±185 MBq for the test-retest study cohort, and 538±150 MBq for the PD cohort. There were no statistically significant differences between the test and retest condition or HS and PD patients in the injected amount of radioactivity, molar activity, or injected mass dose.

### PET imaging experiments

All subjects participated in at least one 60-min [^11^C]UCB-J scans on the High Resolution Research Tomograph (HRRT, Siemens) (17). All participants in this study underwent arterial blood sampling. Radioactive concentration in plasma and parent fractions were measured to generate input function. The details of the procedures of sampling and blood analysis were previously described (6).

In the test-retest study, each subject participated in two [^11^C]UCB-J scans. There were ten same-day and twenty longitudinal (1 week-1month: 4 scan pairs, 1month-1year: 5 scan pairs, >1 year: 11 scan pairs, median 391 days, range: 7-1028 days) scan pairs. For the same-day group, the average time between injections was 5.3 hours (range: 3-7 hours) and they were allowed to consume a light lunch during the interval between the two PET measurements.

### Image analysis

Processing of T1-weighted magnetic resonance (MR) images included skull-and muscle stripping procedures (FMRIB’s Brain Extraction Tool, http://fsl.fmrib.ox.ac.uk/fsl/fslwiki/BET).

The CS2mL and substantia nigra (SN) ROIs were defined in the automated anatomical labeling (AAL) template (18) in our previous work (10, 13, 19). For those ROIs, an average PET image (0–10 min) was aligned to each subject’s MR image via a rigid registration with mutual information. The individual MR image was spatially normalized to the AAL template in the Montreal Neurological Institute space using an affine linear plus nonlinear registration (BioImage Suite 2.5) (20). All other ROIs were defined based on FreeSurfer segmentation on the individual MR images (21). The following FreeSurfer GM ROIs were created by merging original FreeSurfer ROIs. For cortical regions, parcellation was defined in Desikan-Killiany Atlas (22): frontal cortex, parietal cortex, occipital cortex, temporal cortex, anterior cingulate, posterior cingulate, putamen, thalamus, caudate, hippocampus, entorhinal, amygdala. For the WM ROI segmentation, the cerebral WM ROI segmented by FreeSurfer was binarized and smoothed using a 10 mm full width half maximum (FWHM) gaussian filter to generate a WM map (23). Ten differently sized ROIs were generated (0.5, 1, 2, 5, 10, 20, 45, 100, and 200 mL) in individual MR space by applying different thresholds on the smoothed WM map denoted as FreeSurfer-based WM (FBWM) ROIs. The ROI sizes did not vary between subjects.

### PET Quantitative analysis

The one tissue compartment model was applied to estimate regional *V*_T._ DVR values were calculated using the CS2mL, and FBWM ROIs as reference regions. For the test-retest cohort, the reproducibility of *V*_T_ and DVR values, calculated for each evaluated ROI using each reference region, were examined by calculation of the TRV as follows:

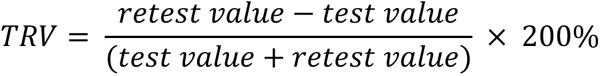

All data are presented as mean ± standard deviation (SD).

For the PD cohort, the percent difference of DVR in the SN, caudate, and brainstem ROIs between HS and PD patient groups were calculated using the optimized FBWM and CS2mL as the reference ROIs. These regions were chosen because they exhibited a significant difference in *BP*_ND_ between HS and PD in Holmes, Honhar et al (21). Statistical analyses between groups were performed with two-tailed, unpaired t-tests, with significance defined as P < 0.05. The between-group percent difference, Cohen’s d, and intersubject %SD using each reference ROI were calculated.

## Results

### V_T_ test-retest variabilities in each reference region

The average *V*_T_ in the CS2mL across each subject’s first scan was 4.33 mL/cm^3^ ± 11.5%. *V*_T_ in FBWM regions increased with increasing ROI size (Fig 1). Only FBWM ROIs 10mL and smaller exhibited less than 10% difference in mean *V*_T_ compared to CS2mL, and %CoV was the second lowest in 10mL (mean values, with %CoVs: 0.5mL: 4.40±13.1%, 10mL: 4.68±10.2%, 20mL: 4.89±10.1%, 100mL: 6.44±12.1%, Fig 1, Supplementary Table S1). While the mean TRV of *V*_T_ in FBWM ROIs are close to 0% and consistent across different region sizes, the TRV SD decreases with increasing region size for both the same-day and longitudinal scan pairs (same-day: 0.5mL: -2.2±14.5, 1mL: -2.4±12.0, 10mL: -1.3±8.5, 20mL: -0.7±8.2, 100mL: 0.5±7.9, and longitudinally: 0.5mL: 0.1±16.9, 1mL: 0.1±17.0, 10mL: -0.5±11.6, 20mL: -0.3±11.2, 100mL: 0.3±10.6, Supplementary Table S2). The *V*_T_ TRV in CS2mL was -0.8±9.9 and -0.6±15.1 for same-day and longitudinal, respectively, demonstrating the higher TRV SD exhibited by CS2mL compared to larger regions such as the 10mL FBWM ROI.

**Fig. 1:**
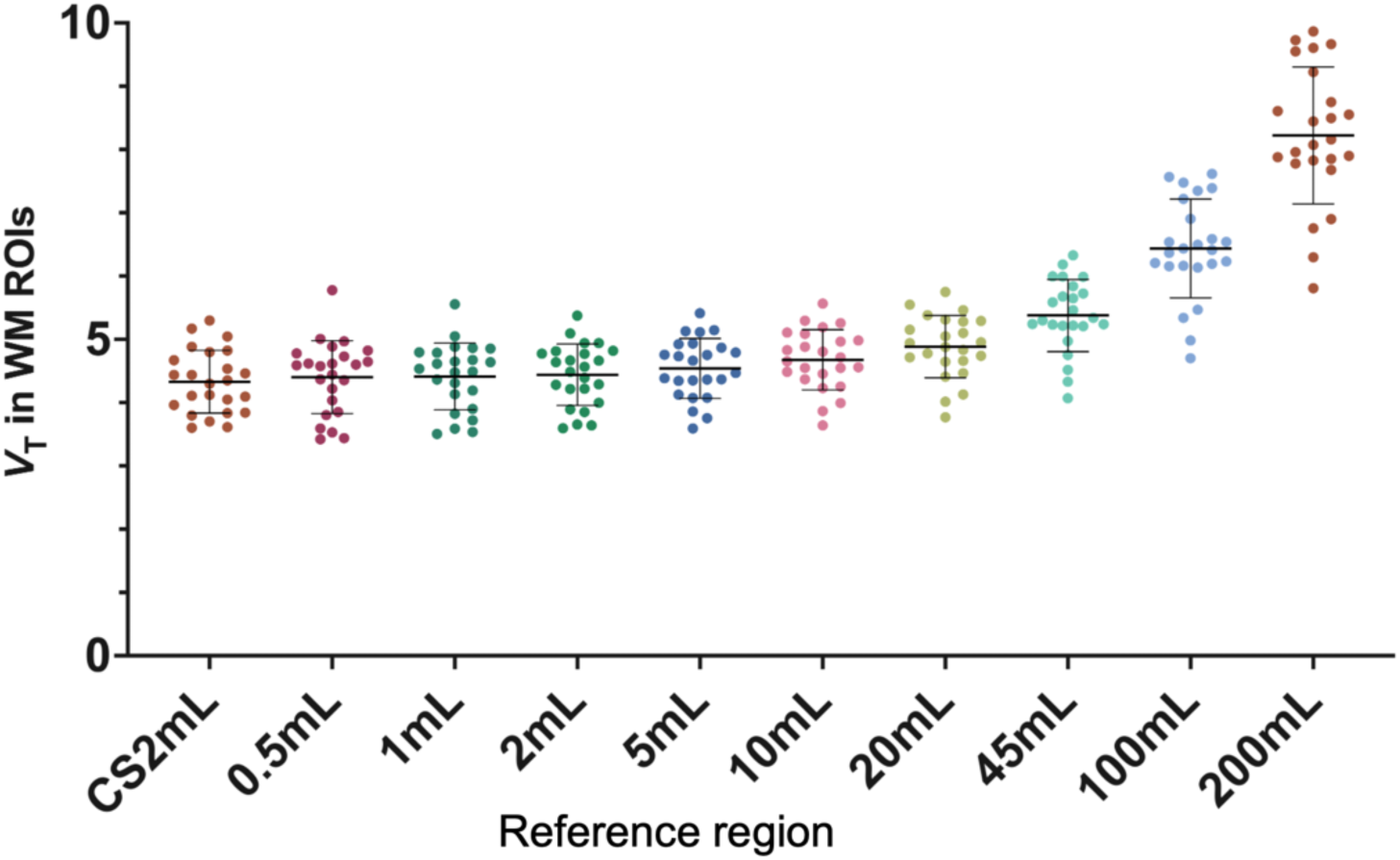
Volume of distribution (*V*_T_) calculated within each reference region for each subject’s first scan (*n* = 24). Regions of interest (ROIs) include the centrum semiovale (CS2mL) and Freesurfer-Based White Matter (FBWM) regions ranging from 0.5-200 mL.

### Test-retest variability in gray matter regions for same-day and longitudinal groups

For each participant in the test-retest study, DVR was calculated for 12 GM regions by dividing the *V*_T_ in each GM region by the *V*_T_ in each reference region. *V*_T_ TRV means and SDs were generally consistent across GM regions. The same-day group exhibited smaller variabilities of *V*_T_ TRV in GM regions compared to the longitudinal group (Table 1). The mean TRV of the DVRs, averaged across all 12 GM regions, were small (*p* > 0.05 for t-test comparing test and retest value) and similar regardless of the reference region suggesting no systematic bias in both same-day and longitudinal group (Supplementary Table S3). The TRV SD of the DVRs, on the other hand, varied based on the reference region used, ranging from 3.1% in the 100mL ROI to 9.8% in the 0.5mL ROI for the same-day dataset, and ranging from 3.8% in the 100mL ROI to 12.4% in the 0.5mL ROI for the longitudinal dataset (Fig 2, Supplementary Table S3). Lower variability in DVR TRV was exhibited by using larger reference regions (Same day: *V*_T_: -0.2±7.6%, CS2mL: -0.9±9.2%, 10mL: 1.2±5.7%, 45mL: -0.3±4.2% 100mL: 0.7±4.4% longitudinal: *V*_T_: 1.0±10.4%, CS2mL: 1.6±11.9%, 10mL: 1.5±7.0%, 45mL: 1.0±5.2% 100mL: 0.7±4.4% table S3). Overall, TRV variability was lower for DVR_2mL-200mL_ compared to DVR _CS2mL_.

**Fig. 2:**
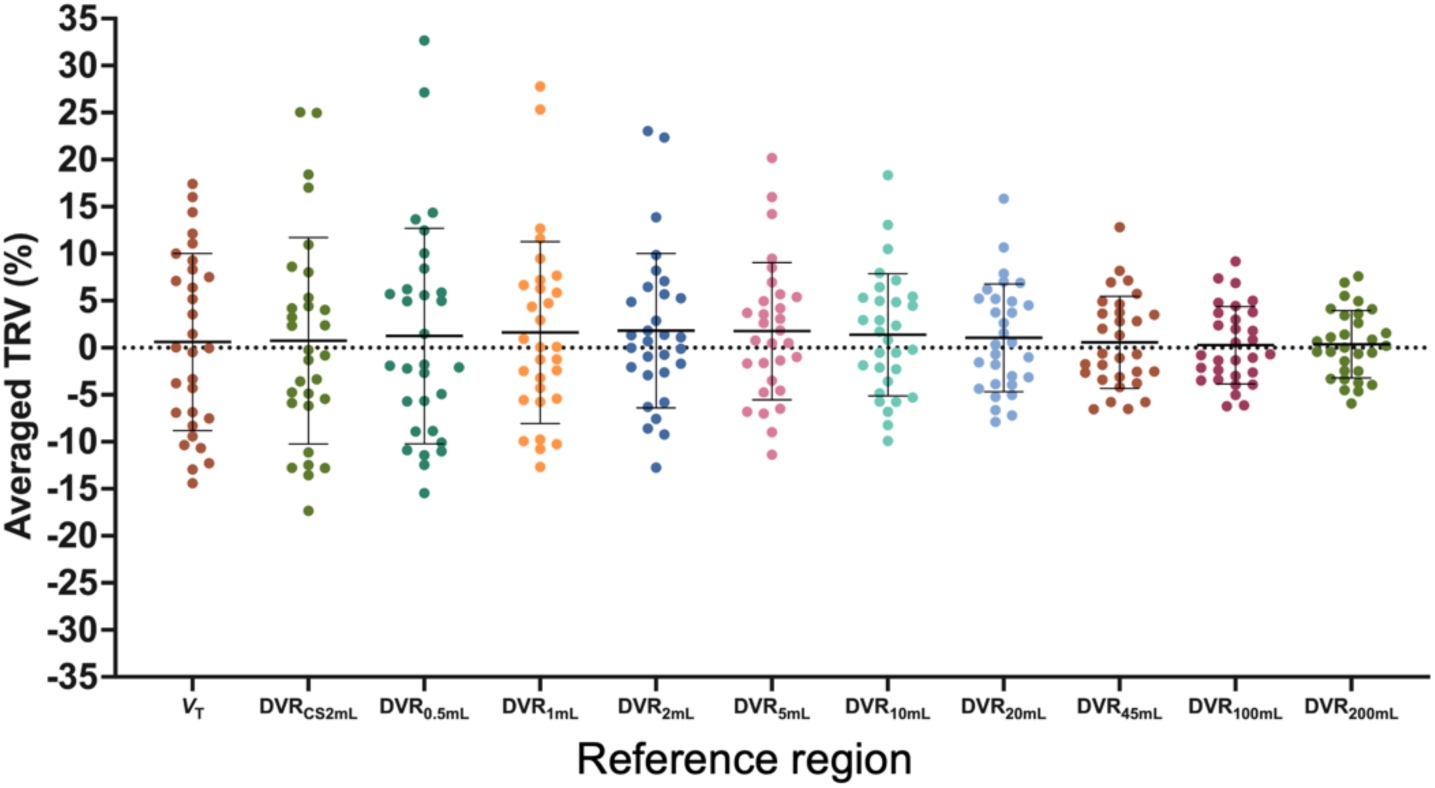
Test-retest variability (TRV) of distribution volume ratios (DVR) calculated using each reference region, averaged among 12 gray matter regions. Reference regions include the centrum semiovale (CS2mL) and Freesurfer-based white matter (FBWM) regions ranging from 0.5-200 mL.

**Table 1:** Same-day and longitudinal test-retest variability (TRV) of volume of distribution (*V*_T_) averaged across subjects for each region of interest.

| | $V_T$ TRV (%) | |
| --- | --- | --- |
|  | Same-day (n=10) | Longitudinal (n=20) |
| Frontal | -0.4±8.3 | 1.3±10.9 |
| Parietal | -0.2±7.1 | 1.4±12.2 |
| Occipital | -0.3±7.6 | 1.7±11.3 |
| Temporal | 0.5±8.5 | 1.4±10.6 |
| Anterior cingulate | 0.5±9.9 | 1.0±11.3 |
| Posterior cingulate | -2.3±7.4 | 1.3±10.6 |
| Putamen | 1.1±8.3 | 0.7±10.0 |
| Thalamus | -3.5±7.7 | 0.2±11.8 |
| Caudate | 1.7±7.4 | 2.0±12.0 |
| Hippocampus | -0.9±8.6 | -0.4±11.3 |
| Entorhinal | 2.4±9.5 | -0.2±9.3 |
| Amygdala | -0.9±6.9 | 0.8±10.9 |

For the longitudinal group, the averaged TRV among 12 GM ROIs were plotted versus the number of days between the first and second scans. For each reference region, the TRV did not significantly change within our study duration, indicating that the inter-scan interval had no consistent effect on TRV up to 1028 days (Fig 3 for *V*_T_, DVR_CS2mL_, DVR_10mL_, and DVR_45mL_). The residual sum of squares was computed for each interscan interval plot using each reference region, demonstrating the reduction in SD induced using larger reference regions (DVR_CS2mL_: 2910.4, DVR_10mL_: 1022.6, DVR_45mL_: 535.8). Balancing the smaller variability achieved using larger ROIs against the small *V*_T_ difference from the previously validated CS2mL region (<10%), and thus small bias, observed in the smaller reference regions, we propose the 10mL FBWM as the optimal reference region.

**Fig. 3:**
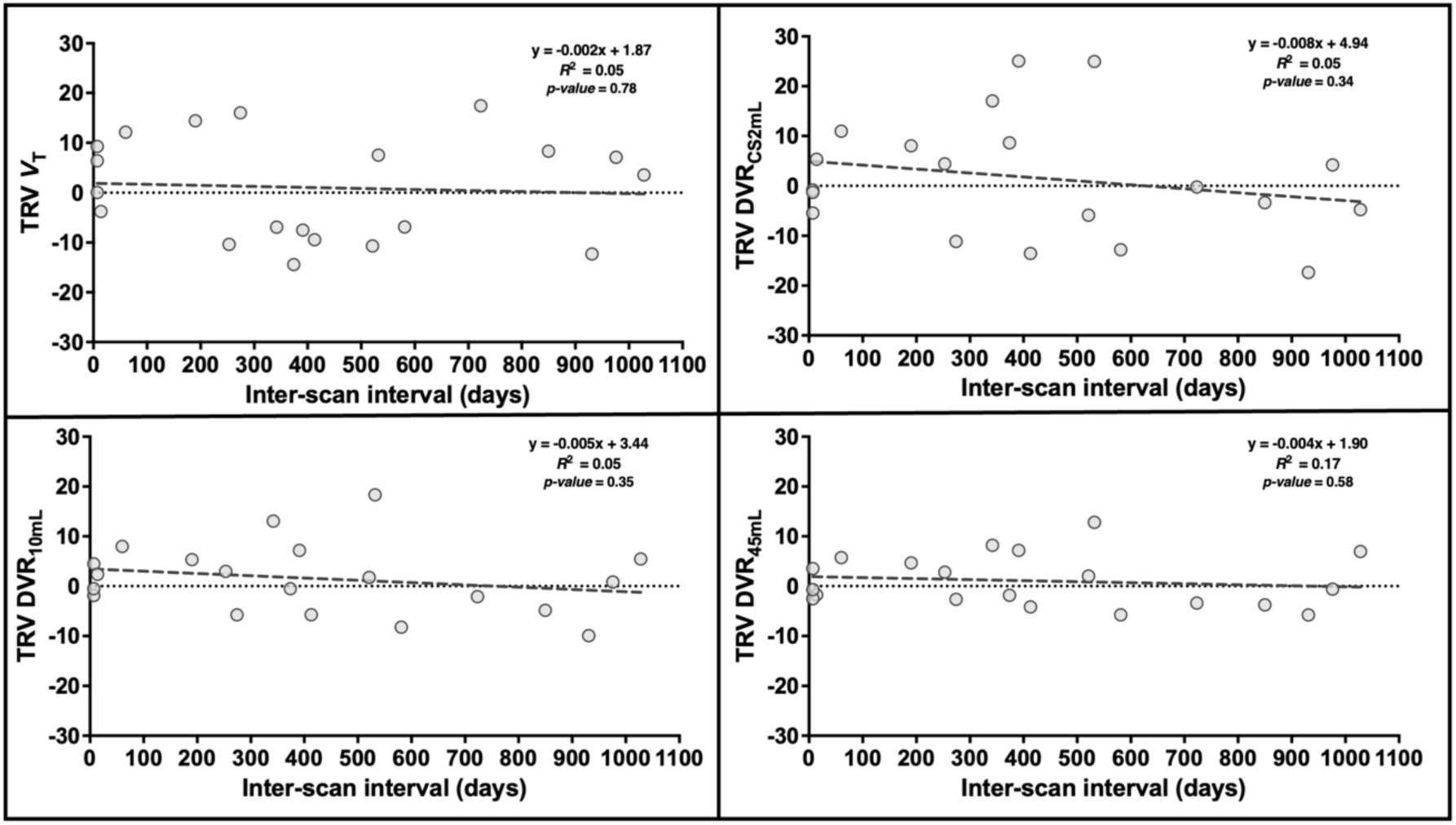
Test-retest variability (TRV) of volume of distribution (*V*_T_) and distribution volume ratios (DVR) calculated using each reference region, averaged among gray matter regions, plotted against interscan interval. Reference regions include the centrum semiovale (CS2mL) and the 10mL and 45mL Freesurfer-Based White Matter (FBWM) regions. The equation of the line of best fit, R^2^, and *p*-values are also displayed for each method.

### FBWM reference region method applied to Parkinson’s disease dataset

For the 10mL reference region, DVRs in the SN and caudate were significantly different between HS and PD groups (p<0.05) but were not significant with the CS2mL reference region (Fig 4). The DVR in the brainstem was not significantly different between groups for either reference region. The intragroup SD for the SN was 16.1% and 14.7% using DVR_CS2mL_ and DVR_10mL_, respectively, for the HS group, while the SD was 18.6% and 16.0% using DVR_CS2mL_ and DVR_10mL_ respectively, for the PD group. The intragroup SD for the caudate was 12.0% and 10.6% using DVR_CS2mL_ and DVR_10mL_, respectively, for the HS group, while the SD was 15.3% and 14.8% using DVR_CS2mL_ and DVR_10mL_ respectively, for the PD group. Further, DVR_10mL_ exhibited a higher group percent difference and Cohen’s d for each ROI, compared to DVR_CS2mL_ (SN: DVR_CS2mL_: 6.5% difference, Cohen’s d: 0.38, DVR_10mL_: 9.0% difference, Cohen’s d: 0.61. caudate: DVR_CS2mL_: 5.7% difference, Cohen’s d: 0.43, DVR_10mL_: 8.0% difference, Cohen’s d: 0.66. brainstem: DVR_CS2mL_: 2.4% difference, Cohen’s d: 0.23, DVR_10mL_: 5.0% difference, Cohen’s d: 0.56. Fig 4, Supplementary Table S4). Of note, the %CoV for *V*_T_ computed in the CS2mL region was 15% and 12% for HS and PD, respectively; for the 10mL ROI *V*_T_ CoV was 11% for both groups (Supplementary Fig S2).

**Fig. 4:**
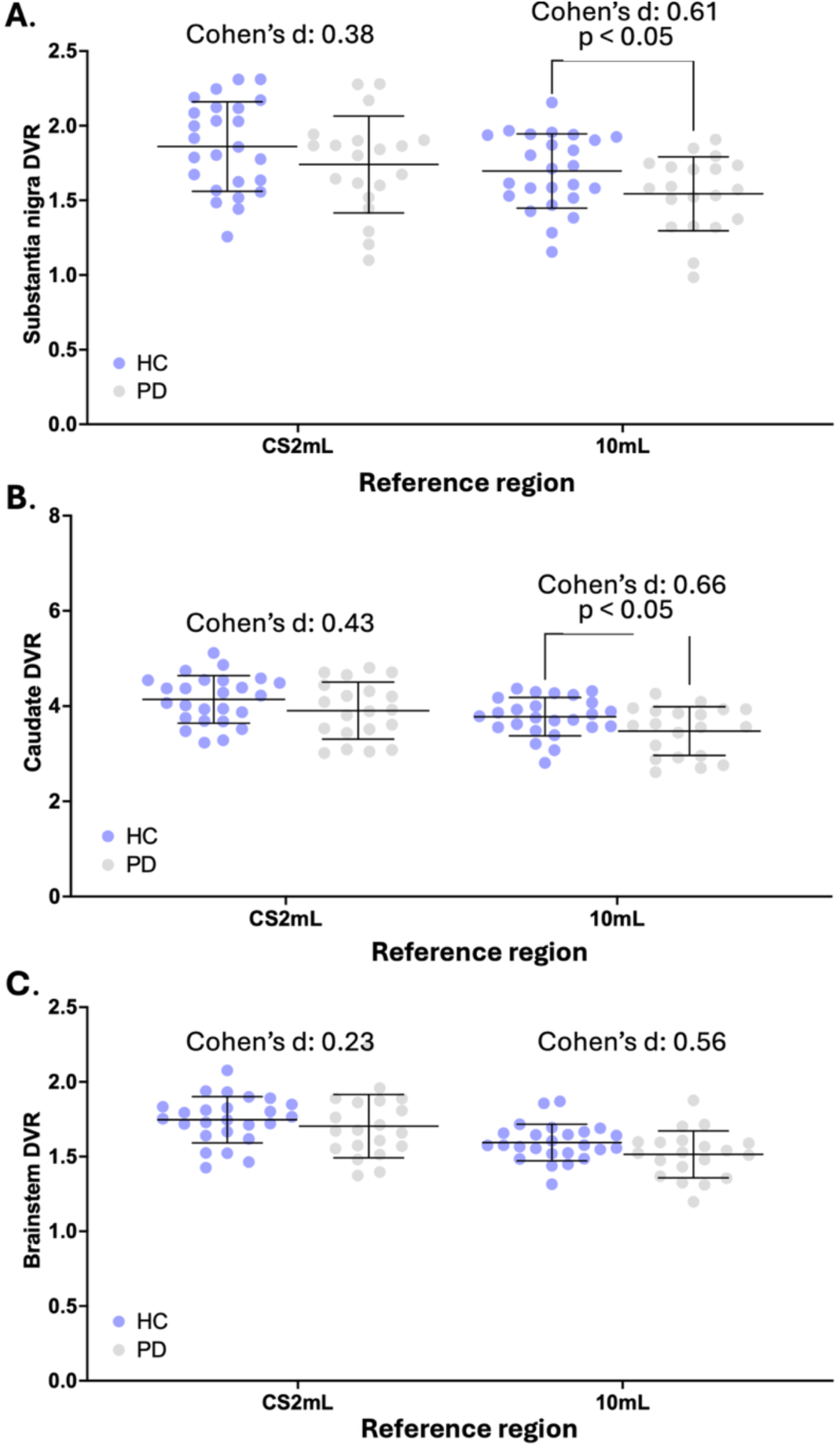
(A) Substantia nigra, (B) caudate, and (C) brainstem distribution volume ratios (DVR) calculated for Parkinson’s disease (PD) and healthy subjects (HS). Reference regions include the centrum semiovale (CS2mL) and 10mL Freesurfer-based white matter (FBWM) region. *p*-values are displayed demonstrating significant difference between HS and PD groups (*p-*value < 0.05). Effect size is displayed using Cohen’s d.

## Discussion

In this study, the longitudinal and same-day test–retest reproducibility using [^11^C]UCB-J was examined, and the development of WM-based reference regions was proposed and applied toward the test-retest reproducibility outcomes and a dataset comparing [^11^C]UCB-J uptake in PD and HS. Overall, we demonstrated that [^11^C]UCB-J exhibits excellent longitudinal test-retest reliability regardless of the reference region used; however, an optimized 10mL WM region can yield [^11^C]UCB-J DVR measurements with reduced inter-individual variation compared to the CS2mL because of its larger size, without introducing significant specific activity (i.e. the 10 mL FBWM ROI was the largest region to exhibit a <10% difference in *V*_T_ compared to the CS2mL, while maintaining the second lowest %CoV across different region sizes). Indeed, a significant group difference between PD and HS groups was elicited from SN and caudate DVR values using the 10mL FBWM region instead of CS2mL. Consequentially, we demonstrated the utility of improved reference regions for [^11^C]UCB-J PET for assessing synaptic density in disorders that necessitate longitudinally reliable measurements.

### Same day and longitudinal test-retest variability

In agreement with our previous study, the same-day TRV in *V*_T_ and DVR for several regions was low, regardless of reference region (6). Similarly, the longitudinal TRVs of regional *V*_T_ and DVR values demonstrated reasonable test-retest reliability, but with higher variability than same-day TRV, sometimes greater than 10%. This is in agreement with a study by Tuncel et al., in which the 28-day test-retest repeatability of [^11^C]UCB-J PET was evaluated, demonstrating a TRV variability of *V*_T_ ranging from 4.2-6.8% (24). Interestingly, using a larger CS region (mean volume 5.4mL), they did not demonstrate a drastic improvement in TRV variability.

There are several potential sources of the higher variability in TRV exhibited in the longitudinal study. First, not all subjects underwent MR imaging on the same scanner, introducing potential variability because MR images are essential for accurate ROI segmentation (25). Further, physiological variability could be introduced by a difference in synaptic density exhibited by some participants between their first and second scan. To this point, only HS were evaluated, with an average age of 39.8 years and longest interval between scans of < 3 years; thus, any change in synaptic density would be unexpected (26). Age-related reductions in synaptic density have been demonstrated in HS (27), and studies have suggested that the caudate exhibits the largest synaptic density reduction per year (28, 29). This pattern is not replicated in our data. To assess longitudinal age-related reductions in synaptic density, an increased sample size and longer interscan interval would be necessary. Nevertheless, an undiagnosed neurological condition in any of the evaluated subjects could result in a change of synaptic density between the scans, introducing inter-individual variability in TRV values. Finally, human- and instrument-induced variability can be associated with several daily processes including radiosynthesis, tracer injection, blood sampling, and scanner calibration (30, 31). This variability likely creates bias throughout the image, providing motivation for using the DVR as the primary measure of specifically bound tracer.

The CS2mL has been optimized and validated as an appropriate reference region with minimum specific binding for calculating [^11^C]UCB-J DVR as well as *BP*_ND_ for some populations (5, 13, 32). Indeed, the CS2mL has been used as a reference region for several studies evaluating regional synaptic density in health (27) and disease (9, 24, 33–35). Although the small size (2 mL in template space, < 2 mL in subject space) renders the previously proposed CS2mL ROI relatively invulnerable to GM spill-in effects that would bias DVR values, it also results in *V*_T_ measurements that are computed from a small number of voxels, increasing DVR variability. Fig 1 visually demonstrates the high variability of the CS2mL *V*_T_, which is propagated into the average DVR_CS2mL_ displayed in fig 3. This phenomenon results in the higher variability in DVR_CS2mL_ TRVs (Table 2) compared to those calculated using any other evaluated reference region.

**Table 2:** Same-day and longitudinal test-retest variability (TRV) of distribution volume ratios (DVRs) calculated using the centrum semiovale (DVR_CS2mL_), 10 mL region (DVR_10mL_), and cerebellum (DVR_CB_) as the reference region, averaged across subjects for each region of interest.

|  | TRV (%) |  |  |  |  |  |  |  |
| --- | --- | --- | --- | --- | --- | --- | --- | --- |
| | $\text{DVR}_{\text{CS2mL}}$ | | $\text{DVR}_{10\text{mL}}$ | | $\text{DVR}_{45\text{mL}}$ | | $\text{DVR}_{100\text{mL}}$ | |
|  | Same-day | Longitudinal | Same-day | Longitudinal | Same-day | Longitudinal | Same-day | Longitudinal |
| Frontal | -1.2±9.5 | 1.9±13.3 | 0.9±6.1 | 1.8±8.5 | -0.6±4.6 | 1.4±6.3 | -0.9±3.9 | 1.1±5.2 |
| Parietal | -0.6±9.5 | 2.0±13.3 | 1.5±5.9 | 1.9±8.8 | 0.0±4.4 | 1.5±6.6 | -0.3±3.6 | 1.2±5.6 |
| Occipital | -0.4±8.7 | 2.3±12.6 | 1.7±4.7 | 2.2±8.2 | 0.2±3.2 | 1.8±6.3 | -0.2±2.4 | 1.5±5.1 |
| Temporal | -0.3±9.4 | 2.0±12.4 | 1.7±5.9 | 2.0±7.2 | 0.3±4.7 | 1.4±5.4 | 0.0±3.9 | 1.1±4.4 |
| Anterior cingulate | -0.3±10.5 | 1.6±13.5 | 1.8±7.4 | 1.5±9.1 | 0.3±6.3 | 1.0±7.2 | 0.7±6.2 | 0.0±5.8 |
| Posterior cingulate | -3.1±8.8 | 1.9±11.3 | -1.0±4.6 | 1.8±7.0 | -2.5±3.1 | 1.3±5.6 | -2.8±2.1 | 1.0±5.1 |
| Putamen | 0.3±8.1 | 1.3±12.4 | 2.4±5.0 | 1.2±7.2 | 0.9±3.9 | 0.7±5.5 | 0.5±3.5 | 0.4±4.8 |
| Thalamus | -4.3±9.4 | 0.8±12.8 | -2.3±5.7 | 0.9±8.0 | -3.7±4.3 | 0.3±6.5 | -4.0±3.4 | 0.0±6.0 |
| Caudate | 1.0±10.8 | 2.6±12.2 | 3.1±7.9 | 2.6±8.1 | 1.6±6.6 | 2.1±7.0 | 1.2±5.7 | 1.8±7.1 |
| Hippocampus | -1.6±9.3 | 0.2±11.4 | 0.5±6.0 | 0.1±7.2 | -1.0±5.1 | -0.3±5.9 | -1.4±4.8 | -0.6±5.6 |
| Entorhinal | 1.6±10.5 | 0.4±11.4 | 3.7±7.7 | 0.3±6.6 | 2.2±7.1 | -0.1±5.8 | 1.9±6.8 | -0.4±5.7 |
| Amygdala | -1.7±9.8 | 1.4±12.1 | 0.4±7.4 | 1.3±8.2 | -1.0±6.3 | 0.9±7.8 | -1.4±5.7 | 0.5±7.6 |

|  | TRV (%) |  |  |  |  |  |  |  |
| --- | --- | --- | --- | --- | --- | --- | --- | --- |
| | $DVR_{CS2mL}$ | | $DVR_{10mL}$ | | $DVR_{45mL}$ | | $DVR_{100mL}$ | |
|  | Same-day | Longitudinal | Same-day | Longitudinal | Same-day | Longitudinal | Same-day | Longitudinal |
| Frontal | -1.2±9.5 | 1.9±13.3 | 0.9±6.1 | 1.8±8.5 | -0.6±4.6 | 1.4±6.3 | -0.9±3.9 | 1.1±5.2 |
| Parietal | -0.6±9.5 | 2.0±13.3 | 1.5±5.9 | 1.9±8.8 | 0.0±4.4 | 1.5±6.6 | -0.3±3.6 | 1.2±5.6 |
| Occipital | -0.4±8.7 | 2.3±12.6 | 1.7±4.7 | 2.2±8.2 | 0.2±3.2 | 1.8±6.3 | -0.2±2.4 | 1.5±5.1 |
| Temporal | -0.3±9.4 | 2.0±12.4 | 1.7±5.9 | 2.0±7.2 | 0.3±4.7 | 1.4±5.4 | 0.0±3.9 | 1.1±4.4 |
| Anterior cingulate | -0.3±10.5 | 1.6±13.5 | 1.8±7.4 | 1.5±9.1 | 0.3±6.3 | 1.0±7.2 | 0.7±6.2 | 0.0±5.8 |
| Posterior cingulate | -3.1±8.8 | 1.9±11.3 | -1.0±4.6 | 1.8±7.0 | -2.5±3.1 | 1.3±5.6 | -2.8±2.1 | 1.0±5.1 |
| Putamen | 0.3±8.1 | 1.3±12.4 | 2.4±5.0 | 1.2±7.2 | 0.9±3.9 | 0.7±5.5 | 0.5±3.5 | 0.4±4.8 |
| Thalamus | -4.3±9.4 | 0.8±12.8 | -2.3±5.7 | 0.9±8.0 | -3.7±4.3 | 0.3±6.5 | -4.0±3.4 | 0.0±6.0 |
| Caudate | 1.0±10.8 | 2.6±12.2 | 3.1±7.9 | 2.6±8.1 | 1.6±6.6 | 2.1±7.0 | 1.2±5.7 | 1.8±7.1 |
| Hippocampus | -1.6±9.3 | 0.2±11.4 | 0.5±6.0 | 0.1±7.2 | -1.0±5.1 | -0.3±5.9 | -1.4±4.8 | -0.6±5.6 |
| Entorhinal | 1.6±10.5 | 0.4±11.4 | 3.7±7.7 | 0.3±6.6 | 2.2±7.1 | -0.1±5.8 | 1.9±6.8 | -0.4±5.7 |
| Amygdala | -1.7±9.8 | 1.4±12.1 | 0.4±7.4 | 1.3±8.2 | -1.0±6.3 | 0.9±7.8 | -1.4±5.7 | 0.5±7.6 |

Thus, a FBWM ROI is likely to be the appropriate choice – that is, if the ROI size is large enough to overcome the variability seen in smaller reference regions but small enough to avoid GM spill-in. Fig 1 and Supplementary Table 1 demonstrate the higher *V*_T_ in larger regions, compared to the CS2mL, likely due to the incorporation of GM voxels containing variable concentrations of [^11^C]UCB-J. On the other hand, the variability in DVR TRVs exhibited by using ≤ 5mL FBWM ROIs (Fig. 2 DVR_0.5-5mL_) was higher compared to FBWM ROIs ≥ 10mL in volume. Tables 1 and 2 demonstrate that longitudinal and same-day DVR_10mL_ TRVs show reduced variability compared to DVR_CS2mL._ These results are in agreement with results published by Tuncel et al; they computed DVRs using a larger CS2mL reference region with a mean volume of 5.4 mL and did not demonstrate a resulting increase in TRV variability. Based on these factors, the 10mL FBWM region was selected for further evaluation using the PD cohort.

### Clinical application: Comparing SV2A binding in PD and healthy subjects

Variability in outcome measurements imposes a limit on the detection of a group difference in [^11^C]UCB-J binding. Studies have demonstrated reduced regional synaptic density in PD subjects compared to HS using the CS as a reference region (9, 10, 36–38). We investigated the impact of using the 10mL FBWM reference region, in place of the CS2mL, to evaluate and compare the variability and group difference in synaptic density measurements within the SN, caudate, and brainstem of PD and HS; these regions were chosen due to their significance, prior to multiple comparisons correction, in another study by Holmes, Honhar et al (9, 10).

The group percent difference for reference region *V_T_* values increased with larger ROI size, with percent differences greater than or equal to 8% only exhibited by ROIs greater than 45 mL in volume (Supplementary Fig. 1). Compared to the CS2mL, *V*_T_ calculated within the 10 mL FBWM ROI exhibited a smaller difference between PD patients and HS, suggesting that the 10 mL ROI does not introduce displaceable binding into the denominator of DVR calculations. Compared to values calculated using the CS2mL, DVR_10mL_ exhibited lower variability, as well as a large percent difference and effect size between the PD and HC groups for each evaluated region. These results are not in agreement with a study by Holmes,

Honhar et. al, in which they found a significant difference in SN, caudate, and brainstem using the CS as a reference region for computing a binding potential (*BP*_ND_) with the simplified reference tissue model 2 method; however, this is likely due to differences in methodology and cohort, the latter of which was larger, including subjects without arterial blood sampling. Of note, several PD subjects included in Holmes, Honhar et al with extensive disease burden (>6 years since diagnosis) did not have arterial blood sampling. Nevertheless, the results herein suggest that the 10 mL FBWM ROI is a better and possibly optimal reference region for studying synaptic density changes in neurodegenerative diseases in which the cerebellum is implicated in disease pathogenesis.

In fact, knowledge of the variability induced by using a particular reference region may inform future study designs. For example, based on a simple power analysis, about 10 subjects are needed to measure a difference, or change, of 10% in synaptic density in the SN, based on the variability measured using the 10mL reference region to calculate DVR in the PD and HS cohorts. This is in comparison to about 15 subjects needed using the CS2mL.

Although the low specific binding of [^11^C]UCB-J and relatively low variability within the 10 mL FBWM ROI encourages its use as a reference region for longitudinal studies, this method is not suitable for use in patients with WM pathology. WM lesions or atrophy, often induced by inflammation such as in multiple sclerosis and HIV encephalopathy, stroke sequelae, and other conditions, can render reference region measurements inaccurate (39). Thus, future optimization of this method should be aimed toward avoiding the inclusion of abnormal WM in reference regions.

In this study we demonstrated the long-term reliability of [^11^C]UCB-J PET in conjunction with new optimized WM reference regions for assessing regional synaptic density in age-related neurogenerative disorders.

## Acknowledgements

The authors appreciate the excellent technical assistance of the staff at the Yale University PET Center. This work was supported by NINDS (R01NS124819; R01NS094253), NIA (R01AG052560), Nancy Taylor Foundation, AbbVie Inc.

## Author contributions

TT, REC, NK collectively contributed to the conception of the study and its design. JDG, MN, REC, and TT contributed to image processing algorithm development. DM, SEH, FES, MM, IE, CHVD, APM, DCD, REC, SJF and TT were responsible for human data acquisition, recruitment, evaluation, and care. YH and NBN were responsible for tracer synthesis and quality control tests. TT and NK performed data analysis. TT, REC, NK, SJF and DM contributed to data interpretation. The first draft of the manuscript was written by NK and TT. All authors participated in the editing the manuscript and approved the manuscript and this submission.

## Statements and Declarations

### Ethical considerations

All procedures performed in studies involving human participants were conducted in accordance with the ethical standards of the institutional and/or national research committee, as well as the 1964 Helsinki declaration and its subsequent amendments or comparable ethical standards.

### Consent to participate

Informed consent was obtained from all individual participants included in the study.

### Consent for publication

Not applicable

### Data availability

The datasets generated during and/or analyzed during the current study are available from the corresponding author on reasonable request.

## Supplementary information

Supplemental material for this article is available online.

