## Supplementary Information for "Optimized reference region and the effect on test-retest reliability and detection of Parkinson’s disease with [^11^C]UCB-J"

**Affiliations:** *[1]Department of Radiology and Biomedical Imaging, Yale University School of Medicine, New Haven, CT, USA; [2]Department of Psychiatry, Yale University School of Medicine, New Haven, CT, USA; [3]Department of Neurology, Yale University School of Medicine, New Haven, CT, USA; [4]Alzheimer's Disease Research Unit, Yale School of Medicine, New Haven, CT, USA; [5]AbbVie Inc., North Chicago, IL, USA*

**Corresponding author:**

Nikkita Khattar, MD-PhD candidate, PET Center, Department of Radiology and Biomedical Imaging, Yale University School of Medicine, New Haven, CT 06520, USA. Tel: +1-203-737-9738, Fax: +1-203-785-3107,

### Supplementary figures

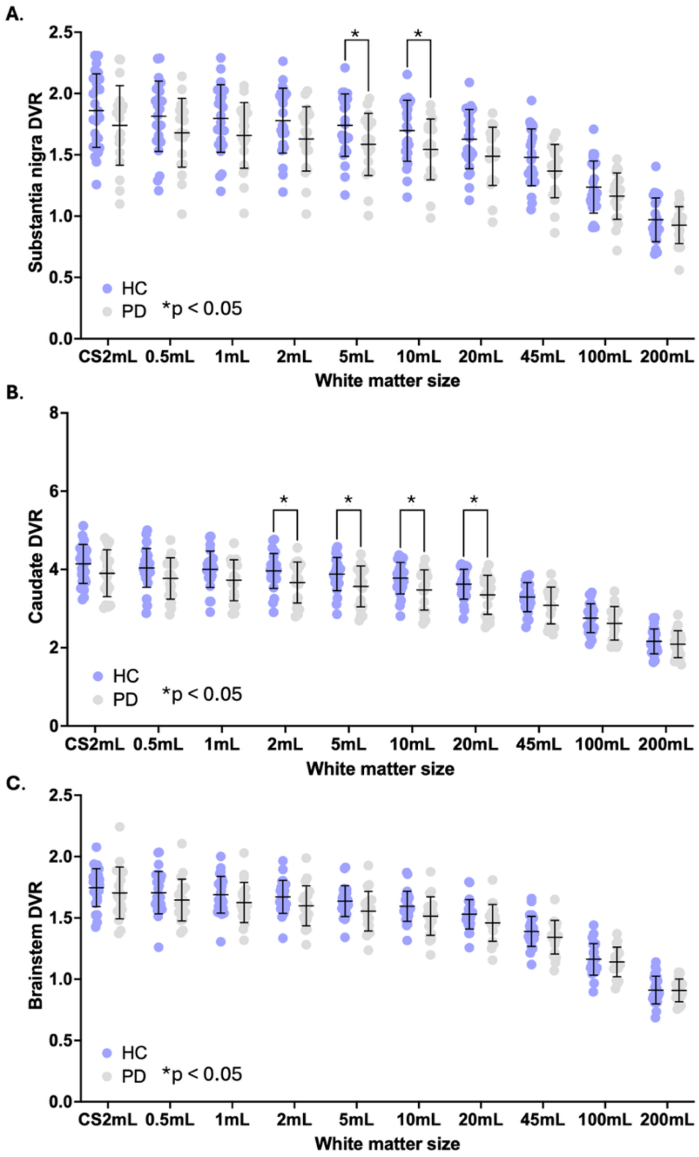


**Fig. S1**: (A) Substantia nigra, (B) caudate, and (C) brainstem distribution volume ratios (DVR) calculated using each reference region for Parkinson's disease (PD) and healthy subjects (HS). Regions of interest (ROIs) include the centrum semiovale (CS2mL) and Freesurfer-based white matter (FBWM) regions ranging from 0.5-200 mL. *p*-values are displayed demonstrating significant difference between HS and PD groups (*$p$-value < 0.05).

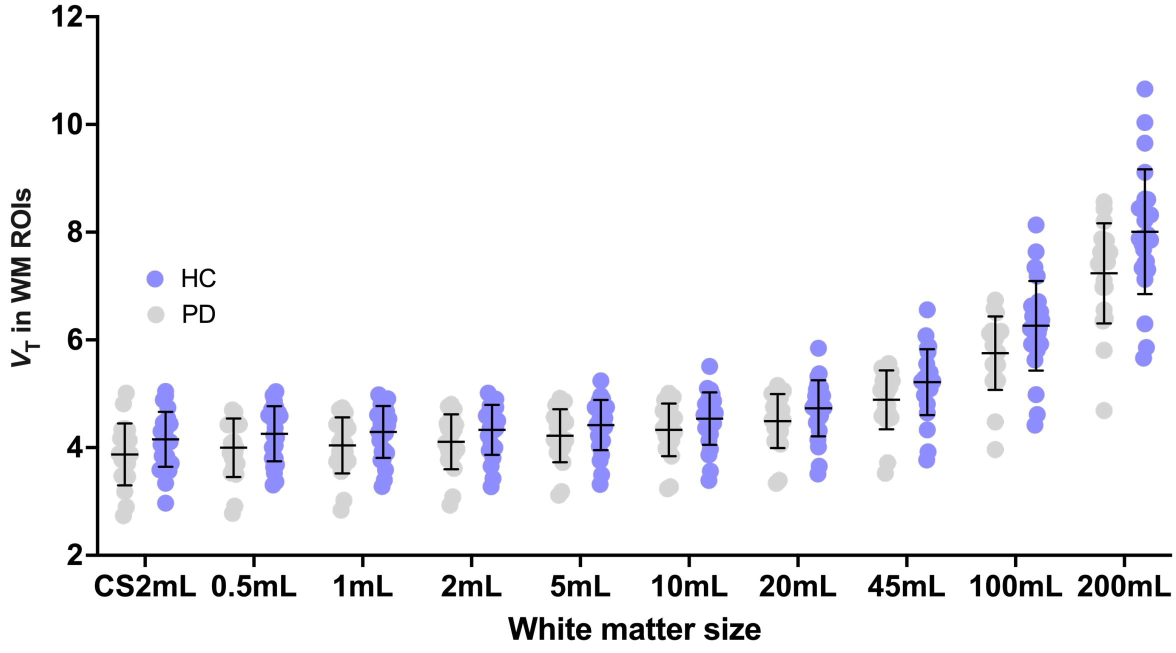


**Fig. S2**: Volume of distribution ($V_T$) calculated within each reference for Parkinson's disease (PD) and healthy subjects (HS). Regions of interest (ROIs) include the centrum semiovale (CS2mL) and Freesurfer-based white matter (FBWM) regions ranging from 0.5-200 mL.

**Supplementary tables:**

**Supplementary Table S1**: Volume of distribution ($V_T$, mL/cm$^3$) calculated for each reference region, averaged across subjects' first scans (n=24) with standard deviation expressed as a percent of mean value. Centrum semiovale is denoted as CS2mL.

| Reference region | $V_T$ |
|---|---|
| CS2mL | 4.33±11.47% |
| 0.5mL | 4.40±13.09% |
| 1mL | 4.41±11.92% |
| 2mL | 4.44±10.93% |
| 5mL | 4.54±10.44% |
| 10mL | 4.68±10.15% |
| 20mL | 4.89±10.07% |
| 45mL | 5.38±10.61% |
| 100mL | 6.44±12.14% |
| 200mL | 8.22±13.16% |

**Supplementary table S2**: Test-retest variability (TRV) of $V_T$ for each evaluated reference region. Values are averaged across subjects.

| Reference region | TRV (%) | |
|---|---|---|
| | Same-day (n=10) | Longitudinal (n=20) |
| CS2mL | -0.8±9.9 | -0.6±15.1 |
| 0.5mL | -2.2±14.5 | 0.1±17.0 |
| 1mL | -2.4±12.0 | -0.3±15.3 |
| 2mL | -2.1±9.7 | -0.7±13.7 |
| 5mL | -1.9±8.7 | -0.8±12.3 |
| 10mL | -1.3±8.5 | -0.5±11.6 |
| 20mL | -0.7±8.2 | -0.3±11.2 |
| 45mL | 0.2±8.1 | -0.9±10.9 |
| 100mL | 0.5±7.9 | 0.3±10.6 |
| 200mL | 0.2±8.0 | 0.3±10.4 |

**Supplementary Table S3**: Test-retest variability (TRV) averaged among gray matter regions, using each reference region in the calculation of the distribution volume ratio (DVR). Mean and SD values across subjects are shown. Centrum semiovale is denoted as CS2mL.

| | **Averaged DVR TRV (%)** | |
|---|---|---|
| **Reference region** | **Same-day (n=10)** | **Longitudinal (n=20)** |
| CS2mL | -0.9±9.2 | 1.6±11.9 |
| 0.5mL | 2.0±9.8 | 0.9±12.4 |
| 1mL | 2.2±7.4 | 1.3±10.8 |
| 2mL | 1.9±6.2 | 1.8±9.2 |
| 5mL | 1.7±6.3 | 1.8±7.9 |
| 10mL | 1.2±5.7 | 1.5±7.0 |
| 20mL | 0.5±5.1 | 1.3±6.1 |
| 45mL | -0.3±4.2 | 1.0±5.2 |
| 100mL | -0.7±3.4 | 0.7±4.4 |
| 200mL | -0.4±3.1 | 0.7±3.8 |

**Supplementary Table S4**: Percent difference and Cohen's d calculated between healthy control and Parkinson's disease group DVRs for the substantia nigra (SN), caudate, and brainstem, for each reference region. Centrum semiovale is denoted as CS2mL.

| | SN | | Caudate | | Brainstem | |
|---|---|---|---|---|---|---|
| Reference region | Percent difference | Cohen's d | Percent difference | Cohen's d | Percent difference | Cohen's d |
| CS2mL | 6.5 | 0.38 | 5.7 | 0.43 | 2.4 | 0.23 |
| 0.5mL | 7.4 | 0.48 | 6.7 | 0.53 | 3.5 | 0.35 |
| 1mL | 7.8 | 0.51 | 6.9 | 0.56 | 3.8 | 0.40 |
| 2mL | 8.4 | 0.57 | 7.5 | 0.61 | 4.4 | 0.49 |
| 5mL | 9.0 | 0.62 | 8.1 | 0.66 | 5.0 | 0.57 |
| 10mL | 9.0 | 0.61 | 8.0 | 0.66 | 5.0 | 0.56 |
| 20mL | 8.6 | 0.58 | 7.6 | 0.62 | 4.6 | 0.52 |
| 45mL | 7.6 | 0.50 | 6.4 | 0.50 | 3.5 | 0.37 |
| 100mL | 6.0 | 0.37 | 4.8 | 0.33 | 1.8 | 0.17 |
| 200mL | 4.6 | 0.27 | 3.3 | 0.22 | 0.4 | 0.03 |
